# Robust spatiotemporal organization of mitotic events in mechanically perturbed *C. elegans* embryos

**DOI:** 10.1101/2023.11.03.565455

**Authors:** Vincent Borne, Matthias Weiss

## Abstract

Early embryogenesis of the nematode *Caenorhabditis elegans* progresses in an autonomous fashion within a protective chitin eggshell. Cell division timing and the subsequent, mechanically guided positioning of cells is virtually invariant between individuals, especially before gastrulation. Here, we have challenged this stereotypical developmental program in early stages by mechanically perturbing the embryo, without breaking its eggshell. Compressing embryos to about 2/3 of their unperturbed diameter only resulted in markedly slower cell divisions. In contrast, compressing embryos to half of their native diameter frequently resulted in a loss of cytokinesis, yielding a non-natural syncytium that still allowed for multiple divisions of nuclei. Although the orientation of mitotic axes was strongly altered in the syncytium, key features of division timing and spatial arrangement of nuclei remained surprisingly similar to unperturbed embryos in the first few division cycles. This suggests that few, very robust mechanisms provide a basic and resilient program for safeguarding the early embryogenesis of *C. elegans*.

**STATEMENT OF SIGNIFICANCE:** Early embryogenesis of the nematode *Caenorhabditis elegans* progresses in an autonomous fashion within a protective chitin eggshell. Cell division timing and cell positioning seemingly runs on autopilot, yielding a stereotypical development. Compressive forces, a potential hazard in the nematode’s native habitat, may jeopardize this. We show that compressing embryos to 2/3 of their native diameter results in markedly slower cell divisions but leaves the early embryonic program otherwise intact. Further compression of embryos impairs the formation of new cells while nuclei still divide in a common cytoplasm (’syncytium’) with basic features of division timing and spatial arrangement being surprisingly similar to unperturbed embryos. This suggests that few robust mechanisms provide a basic program for the early embryonic autopilot.

## I. INTRODUCTION

Embryogenesis is a fundamental and complex process that embraces the emergence of a full-grown multicellular organism from a single cell. Starting from a fertilized oocyte, subsequent cell divisions and the spatial arrangements of these newly formed building blocks into specialized tissues need to be timed and directed without external help. Embryos therefore need to self-organize on many length and time scales and even supposedly simple model organisms like the nematode *Caenorhabditis elegans* feature an impressive embryonic self-organization already at early stages. In fact, *C. elegans* embryos determine their body axes during the first three rounds of cell divisions and alongside they also define precursor cells for distinct tissues, e.g. the germline [1]. Moreover, embryos follow a stereotypical development with an invariant cell lineage tree [2] (see Fig. 1a for the first four generations of cells), eventually yielding in 99% of all cases a hermaphrotitic adult animal with 959 somatic nuclei.

**FIG. 1.**
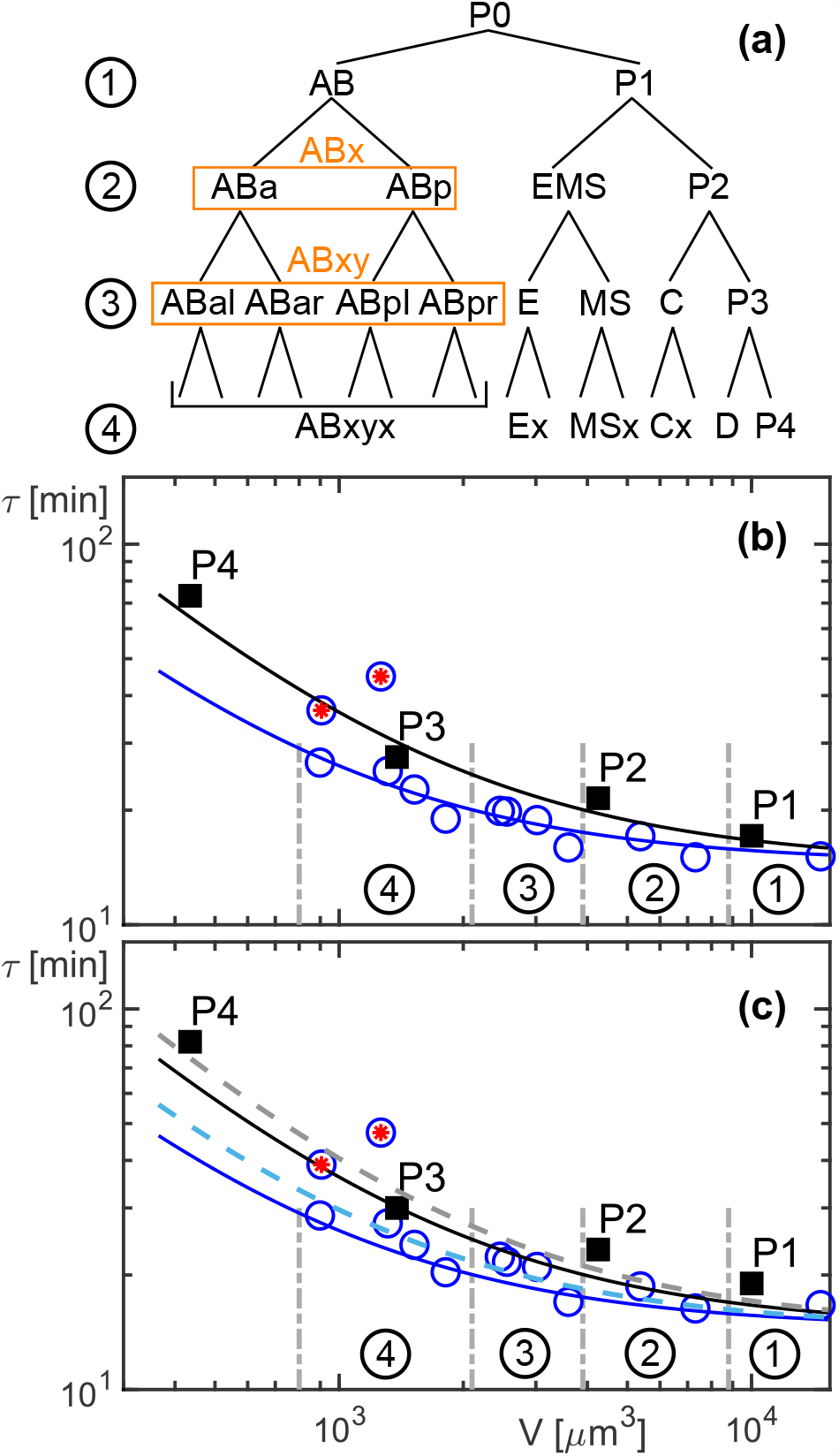
(a) Lineage tree of the early embryo. Cells of generation 1 (AB, P1) produce daughter cells ABa, ABp, EMS, P2 (generation 2). Somatic cells are partially grouped for the analysis (indicated by orange boxes with labels x=a/p and y=l/r). (b) Lifetimes *τ*, i.e. anaphase to anaphase periods, obtained from unperturbed embryos (*n* = 12) are anti-correlated with cell volumes *V*. Somatic cells (open blue circles) and germline cells (filled black squares) follow the previously found powerlaw relation [Eq. (1)] (blue and black lines, refered to as benchmark curves). Vertical dashed lines and encircled numbers indicates the cell generation (note that P3 and P4 belong to generations 3 and 4, respectively). Cells D and Ea/Ep are highlighted by red asterisks. (c) Gentle compression of embryos to a diameter of 20 *μ*m (*n* = 3) yielded slightly larger lifetimes (germline: 10%, somatic: 5-12%) that were still in reasonable agreement with the benchmark curves; same plot style as before. Increasing prefactors *α* in Eq. (1) by 20-30% yielded an improved description of the data (blue and grey dashed lines). See main text for details. Standard errors of the mean are smaller than symbol size in all cases.

Given this striking reproducibility, its optical transparency, and the ease with which genetic changes can be induced, *C. elegans* is a widely used model system for studying self-organisation during (early) embryogenesis. Combinations of advanced light microscopy and theoretical modeling have revealed, for example, detailed physico-chemical insights into the first, asymmetric cell division [3, 4], the onset of cytoplasmic streaming [5–7], and the associated Turing-like pattern formation that supports the definition of germline precursors [8–11]. Moreover, chiral force fields have been seen to govern divisions of somatic cells, eventually also determining the animal’s left-right body axis [12, 13]. In addition, repulsive forces between neighboring cells and/or the engulfing chitin egg shell have been revealed as key determinants for the migration and spatial arrangement of cells until gastrulation [14]. Combining this feature with the observation of an inverse relation of cell volume and cell division times [15, 16] suggests that the early embryogenesis of *C. elegans* is an almost deterministic process that virtually runs on autopilot.

Given that *C. elegans* embryos seem to ‘know’ how to time cell divisions and to spatially arrange the resulting cellular building blocks in a self-organized fashion, we wondered how these fundamental ingredients for successful embryogenesis would change in response to perturbations. In particular, we asked how robust the timing of mitosis events would be maintained and how space compartmentalization might be altered when driving embryos out of their comfort zone by external cues. Previous data have revealed already that temperature changes within the physiological range only lead to an Arrhenius-like rescaling of division times [16], i.e. in cold conditions the entire development simply progresses slower. Only upon exceeding a critical temperature, protein misfolding and degradation has been reported to lead to an aberrant development [17]. As to aspects of spatial organization, an altered but still optimal cell positioning was observed in very early stages of development when forcing *C. elegans* embryos into pre-defined geometries, from circles to prolate ellipsoids [18]. In fact, all observed cell patterns in *C. elegans* embryos and in these artificially constrained geometries are well predicted when modeling cells as soft spheres that mutually interact in a basin with fixed boundary, forcing cells to relax into positions of least mechanical constraints [14, 18].

Here we have followed up on these approaches and explored how a compression of early *C. elegans* embryos, a potential hazard in the native habitat, affects the timing and sequence of mitotic events.

## II. MATERIALS AND METHODS

Embryos from different transgenic *C. elegans* strains were used for this study: In JH2840 (pgl-1::GFP) [19], the p-granule constituent protein PGL-1 is tagged with a green fluorescent protein (GFP); OD95 (GFP::PH(PLC1*δ*1) + mCherry::his58) [20] features a GFP-labeled plasma membrane and mCherry-labeled histones; XA3051 (GFP::H2B + GFP::tbb-2) [21], a kind gift of I. Mattaj (EMBL), expresses GFP-tagged *β*-tubulin and histones. Worms were stored and cultured following standard protocols [22]. Nematode dissection and egg extraction were performed as described previously [23].

Embryos that are refered to as ‘unperturbed’ were placed in 50 *μ*m or 120 *μ*m deep wells (both were seen to not deform the embryo): 50 *μ*m wells were obtained by using double-sided tape as a spacer into which 6 mm diameter holes were punched. The tape was then glued onto a 24 mm×60 mm #1 coverslips (Epredia; Thermo Fisher Scientific, Dreieich, Germany), and single embryos were placed in the hole with a 2 *μ*l droplet of M9 buffer. Holes were sealed by a 20 mm×20 mm #1 coverslip (Menzel; Thermo Fisher Scientific, Dreieich, Germany) that was placed on top of the tape to avoid evaporation of the medium during the recording. For 120 *μ*m wells, SecureSeal™ imaging spacer (Grace Bio-Labs, Bend, OR, USA) was used, following the same protocol.

For compressed embryos, polystyrene beads with diameters 20 *μ*m or 12 *μ*m were used as spacers (see Supplement, Fig. S1a-c): Eggs were placed directly onto #1 coverslips with a droplet of M9 buffer. An additional 2 *μ*l of M9 medium, pre-mixed with a 1% solution of Micromod polystyrene beads (Micromod, Rostock, Germany) was then added and a second coverslip was placed on top, resting on the dispersed polystyrene beads. To avoid movements and medium evaporation, hot Vaseline was carefully applied around the upper coverslip to seal the chamber. Due to bead polydispersity (19.5 − 20.5 *μ*m and 11.5 − 13.5 *μ*m, respectively) and since deformed embryos were expected to exert an elastic force against compression, the actual chamber height was controlled by marking both coverslips with a pencil and using the z-stage controller to check the distance between these two marks by the focus plane of the confocal microscope. Height control prior to and directly after each time series acquisition revealed an average sample thickness of 20 *μ*m and 14 *μ*m, respectively, without a significant drift between the two times of measurement (Fig. S1d).

All measurements were performed on a customized spinning-disk confocal microscope, consisting of a Leica DMI 4000 microscope body (Leica Microsystems, Mannheim, Germany), equipped with a SCAN IM 130×85 sample stage (Maerzhaeuser, Wetzlar, Germany), a CSU-X1 (Yokogawa Microsystems, Tokyo, Japan) spinning disk unit, and a Hamamatsu Orca Flash 4V2.0 sCMOS camera (Hamamatsu Photonics, Hamamatsu City, Japan). The setup was controlled using a custom-made code in LABVIEW (National Instruments, Austin, TX, USA), coupled to the HOKAWO imaging software interface (Hamamatsu Photonics, Hamamatsu City, Japan) for image acquisition. Samples were kept at 22.5°C throughout and were imaged with a HC PL APO 63x/1.4 (Leica Microsystems, Mannheim, Germany) oil immersion objective. Illumination of the specimen at 491 nm and 561 nm was achieved by two solid-state lasers (Calypso and Jive, Cobolt AB, Stockholm, Sweden) in a Dual combiner. The illumination power, measured at the back aperture of the objective, was set to 0.04 mW for 491 nm and to 1.2 mW for 561 nm unless stated otherwise. Fluorescence signals were detected by bandpass filters (Semrock, Rochester, NY, USA) in the range 500-550 nm and 575-625 nm, respectively.

Over a time course of 2-3 h, image stacks of 30 confocal image planes (separated by 1 *μ*m) were acquired every 60 s with 50 ms exposure time for each layer. Recording of a single stack required 8-9 s, i.e. a three-dimensional reconstruction of early embryos was possible. Image series were analyzed using ImageJ/FIJI with the plugin TrackMate [24] and custom-made codes in Matlab (The MathWorks, Natick, MA, USA) for tracking nuclei and cell divisions [15]; all subsequent analyses were performed with custom-made Matlab codes.

## III. RESULTS AND DISCUSSION

Aiming to mechanically perturb the seemingly deterministic early development of *C. elegans*, we have exposed individual embryos to compressive forces by sandwiching them between two coverslips (see Materials and Methods). Since *C. elegans* embryos have an axisymmetric ellipsoidal shape with a long axis of about 50 *μ*m and a diameter of 30 *μ*m, we have used spacings of 50 *μ*m or larger to monitor unperturbed embryos, whereas spacers ≤ 20 *μ*m were used to reduce the diameter of embryos.

As a sensitive readout that reflects on the early developmental program, we have considered the lifetime *τ* of individual cells (i.e. the period between successive anaphases) along the lineage tree for the first four generations (cf. Fig. 1a). For unperturbed embryos, these lifetimes *τ* have been shown to display a clear anti-correlation with cell volume *V* [15, 16], i.e.

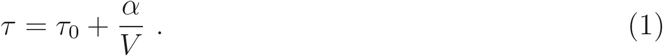

Here, *τ*_0_ ≈ 870 s denotes a cell-independent minimum time that is needed for basic processes like chromatin (de)condensation. The parameter *α* is a constant for somatic and germline cells until gastrulation, but differs about twofold between these two lineages (*α*_*g*_ = 1.86*α*_*s*_ ≈ 2.2×10^4^ *μ*m^3^ min). No changes were seen when altering the size of embryos via RNAi or when releasing internal mechanical constraints by removing the chitin eggshell [15]. Therefore, we will refer to these two lineage-specific relations *τ* (*V*) as ‘benchmark curves’ in the following discussion. Notably, upon varying the ambient temperature in the physiological range, from 15°C to 25°C, all values for *τ* were seen to change according to a common Arrhenius factor [16], i.e. the developmental program was robust in its timing sequence but with an altered overall time scale (slower development at lower temperatures).

All of these results have been revealed with experimental data from lightsheet microscopy, in which illumination-induced phototoxic effects are minimized in comparison to alternative fluorescence imaging techniques [25]. We have therefore probed in a first set of experiments whether also confocal imaging of unperturbed *C. elegans* embryos yields data for *τ* (*V*) that are compatible with the benchmark curves. Since confocal imaging at high resolution is gradually compromised in regions that are more distal to the objective, determining cellular volumes in the 30 *μ*m thick embryo yielded poor results. We have therefore used the previously reported mean cell volumes along the early lineage tree [26] to complement the lifetimes extracted from confocal imaging. As a result, we observed that confocal imaging of unperturbed, i.e. uncompressed, embryos yielded cell lifetimes that were in favorable agreement with the benchmark curves from lightsheet imaging (see Fig. 1b). In the fourth generation, outlier cells D and Ex were seen to deviate from the benchmark curve of somatic cells, featuring significantly larger lifetimes. Such deviations to larger times are actually expected as these cells are the first in which the cell cycle includes an appreciable gap phase whereas the other cells merely cycle between S- and M-phase [27].

As a next step, we gently compressed embryos to a slightly flattened ellipsoid by using 20 *μ*m spacers for the sandwiching process, i.e. the embryonic diameter was reduced by ∼ 10 *μ*m. Such a gentle compression, which leaves the eggshell intact, is widely used as sample mounting technique for confocal imaging of *C. elegans* embryos [23, 28–30] as it reduces the aforementioned limitations on imaging distal regions but allows for live imaging throughout embryogenesis, i.e. the development is not impaired. As a result, we observed that lifetimes in gently compressed embryos were still in reasonable agreement with the benchmark curves although the average values for *τ* were increased by about 10% (Fig. 1c). In particular, the lineage-specific relations *τ* (*V*) were preserved, with lifetimes of germline cells being consistently larger than those of somatic cells with comparable volume. In fact, keeping *τ*_0_ in Eq. (1) constant but increasing *α*_*g*_ by 20% and *α*_*s*_ by 30% (dashed curves in Fig. 1c), yielded an improved description for the experimental data (as judged from the reduced residuals). Altogether, we can conclude from these data that the timing and sequence of mitotic events in the early development of *C. elegans* is very robust and almost unaffected when applying compressive forces that reduce the embryo to about 2/3 of its native diameter.

Interestingly, when challenging embryos with phototoxic stress, induced by an approximately eightfold increased illumination intensity in the blue range (from 0.04 mW to 0.3 mW at 491 nm), we observed severe changes already in the first three generations of cells. For uncompressed embryos, cell lifetimes were on average 30% higher than the benchmark curves (see Fig. 2a) and the clear difference between relations *τ* (*V*) for somatic and germline cells was not visible anymore. Still, the general anti-correlation of *τ* and *V* was preserved. Gently compressed embryos (using again 20 *μ*m spacers) were seen to be even more sensitive to the illumination-induced phototoxicity, suggesting a synergistic effect of both perturbations. All cell lifetimes were significantly higher than the benchmark curves (on average 57%) and the difference between somatic and germline cells was erased (Fig. 2b). Yet, the general anti-correlation of *τ* and *V* according to Eq. (1) was maintained and provided (with a strongly increased value for *α*) a reasonable description of cell lifetimes irrespective of the lineages (cf. red dashed line in Fig. 2b). From these data we can conclude that the general scaling [Eq. (1)] is a very robust feature of early *C. elegans* embryos, even when slowly poisoning the (squeezed) organism via phototoxic effects.

**FIG. 2.**
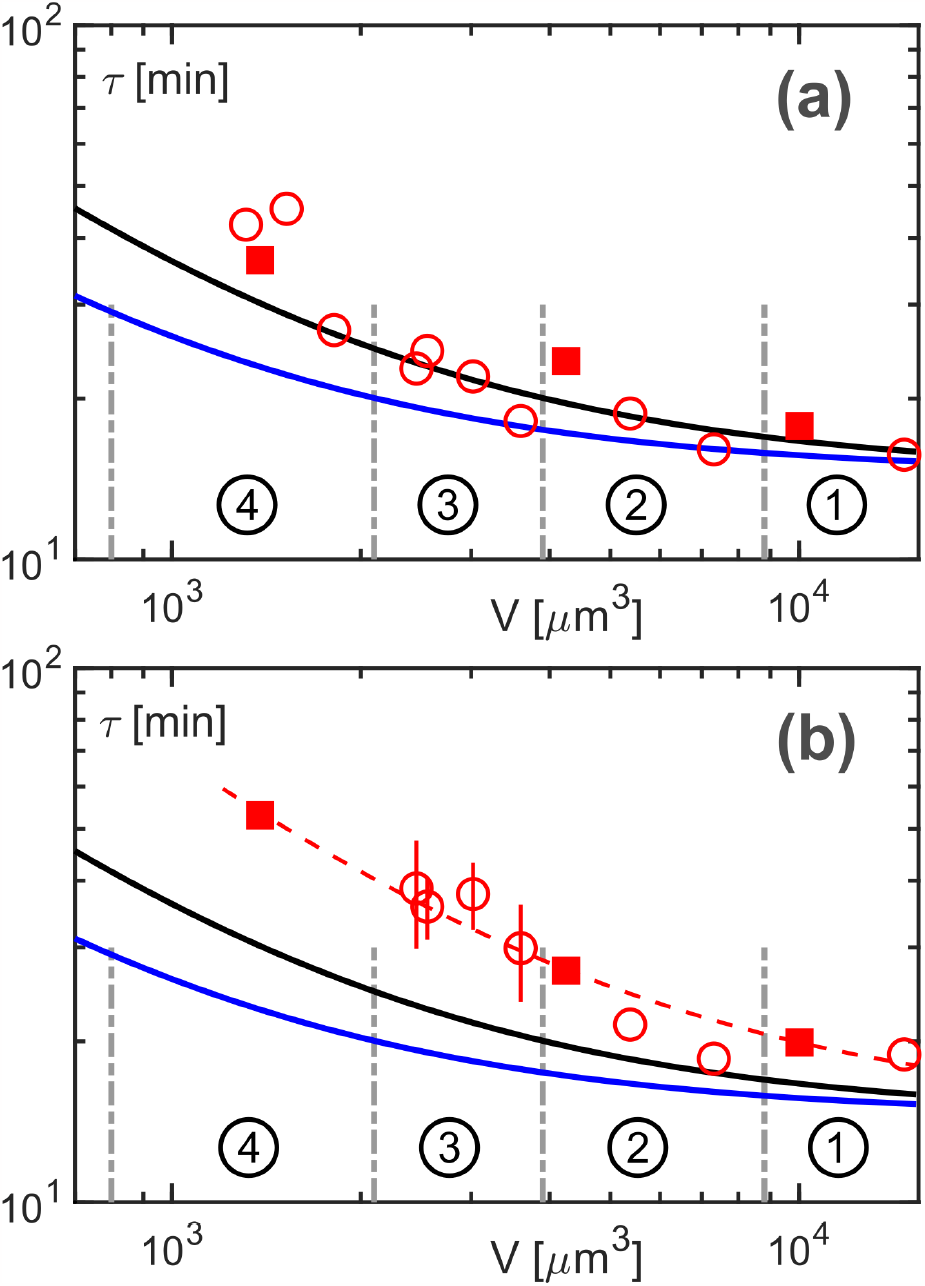
Cell lifetimes *τ* versus volumes *V* for (a) unconstrained embryos (*n* = 28) and (b) gently compressed embryos (thickness 20 *μ*m, *n* = 7), when imaged at a markedly increased illumination power in the blue range (0.3 mW at 491 nm). Somatic and germline cells are shown as open circles and filled squares, respectively. In both cases, clear deviations from the benchmark curves (blue and black lines) are visible, supposedly due to photoxic effects that impede proper embryogenesis. Moreover, data sets for somatic and germline cells now overlap, hence perturbing the chronology of cell division events. Yet, even for the combined challenge of compression and photoxicity, the general anti-correlation of *τ* and *V* according to Eq. (1) is preserved (unaltered *τ*_0_ but *α* = 2.5*α*_*g*_, red dashed line). Error bars (reflecting the standard error) are only shown when larger than symbol size.

As a next step, we increased the mechanical perturbation by using 12 *μ*m spacers to compress *C. elegans* embryos to about half of their native thickness (cf. Supplement, Fig. S1d). Upon doing that, we frequently observed a total failure of cytokinesis already at the first cell division (54% of all experiments with these spacer, but 75% of all experiments in which the embryo attained a diameter ≤ 15 *μ*m). The resulting non-natural syncytium state (similar to the syncytial blastoderm of *Drosophila melanogaster*, cf. [31]) still featured mitotic events and multiple rounds of nucleus duplications (see Fig. 3) that allowed us to define nucleus generations in full analogy to cell generations in unperturbed embryos.

**FIG. 3.**
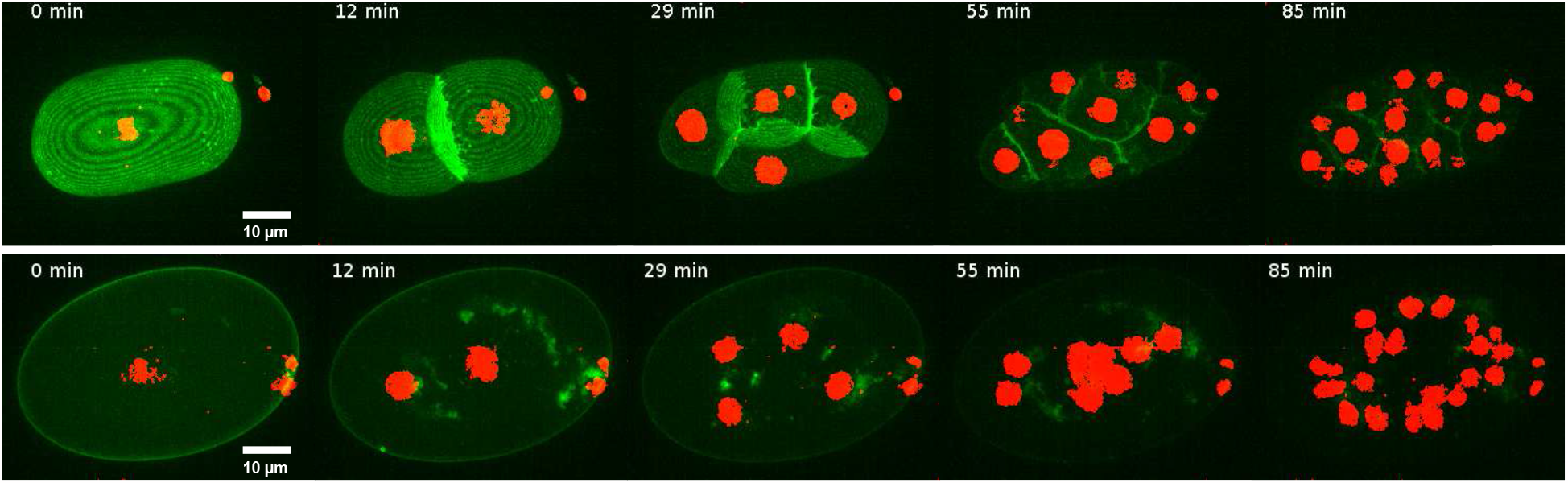
Representative fluorescence image series (z-projections) of the development of an unperturbed embryo (upper panel) and a strongly squeezed embryo in which cytokinesis fails but nuclei can still divide in a syncytium (lower panel). Green fluorescence indicates the plasma membrane of cells while chromatin is highlighted in red.

Lifetimes of syncytial nuclei (defined again as the period between successive anaphases) were still seen to increase with every generation, in accordance with cell lifetimes in unperturbed embryos (Fig. 4). This observation suggests that the early developmental program relies on a robust clock that does not require a compartmentalization by distinct cells. It also highlights that not the cellular volume as such but rather the amount of cytoplasm or the available number of cytoplasmic factors per nucleus are key for an increasing lifetime in successive generations. In line with this notion and in contrast to unperturbed embryos, nuclei in the syncytium were seen to always divide in a synchronous rather than in a sequential fashion. Therefore, the precise timing for sequential divisions within each generation, potentially ensuring a proper relaxation of cells to their desired locus [15], is compromised in the syncytium state. Notably, both observations are in accordance with previous reports on single embryos that have been permeabilized by laser ablation to block cytokinesis via the actin-severing drug cytochalasin D [32].

**FIG. 4.**
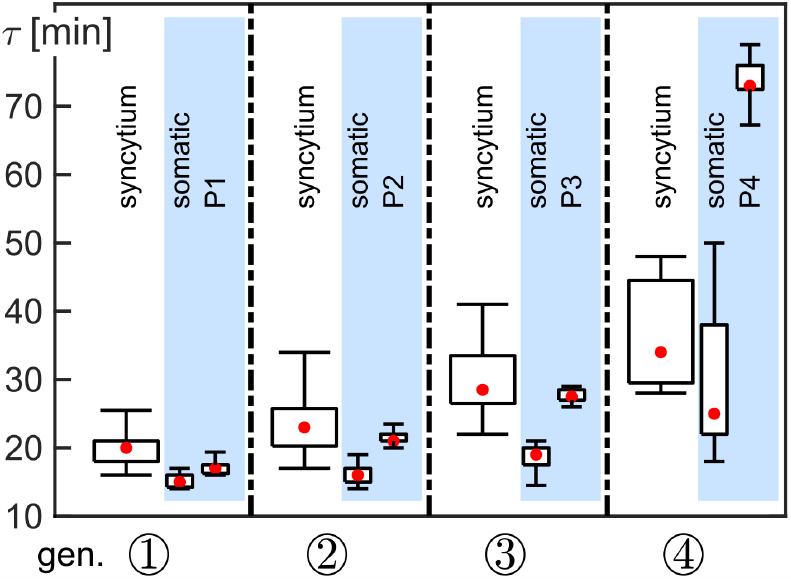
Comparison of lifetimes *τ* of cells/nuclei in unperturbed embryos (light blue background, *n* = 12) and squeezed embryos that have developed a syncytium (white background, *n* = 27). The comparison utilizes nucleus generations for ordering, (2*°* dividing nuclei in generation *n*). In generation 1, *τ* is significantly larger in the syncytium than for the corresponding somatic and germline cells (two-sample Kolmogorov-Smirnov test with threshold *p* = 5%). In generations 2 and 3, values for *τ* are significantly larger in the syncytium than in somatic cells but do not significantly differ from germline cells, P2 and P3. For generation 4, syncytial nuclei feature an intermediate lifetime, between that of somatic and germline cells in unperturbed embryos.

A quantitative and generation-wise comparison of nucleus lifetimes further revealed that the timing of mitotic events in the syncytium is quite close to that of germline precursors in the unperturbed embryo (Fig. 4). This observation is a bit surprising as one might have expected the syncytium to feature lifetimes that are between those seen for somatic and germline cells [32]. To rationalize this finding, we recall that p-granules are the defining features of germline precursors, i.e. they are only observed in P cells. Given that P cells feature lifetimes that are significantly longer than those of equally sized somatic cells (cf. Fig. 1b and Ref. [15]), it is reasonable to assume that p-granules are invoked in prolonging the cell cycle time, e.g. by sequestering mitosis-promoting factors. Interestingly, the syncytium state still allows for the assembly and gradient formation of p-granules (see Supplement, Fig. S2), i.e. p-granules should also able to prolong (and maybe even to synchronize) the cycle time for the division of syncytial nuclei. In fact, p-granules locate initially to the posterior pole of the syncytium (Supplement, Fig. S2c), similar to what is seen in unperturbed embryos before the first cell division [33]. At later stages, p-granules rather assume a perinuclear localization pattern in the syncytium (Supplement, Fig. S2d), akin to what is seen in P4 cells of unperturbed embryos. Based on these observations, one can actually expect the divisions of nuclei in the syncytium to follow the timing of germline precursors for the first generations due to the presence of p-granules near to the posterior pole. In the fourth generation, in which the syncytial division timing rather approaches the timing of somatic cells (cf. Fig. 4), the perinuclear arrangement of p-granules might not be able anymore to prolong the cell cycle markedly.

Altogether, the observed timing of mitosis events in the non-natural syncytium state is remarkably close to what is seen in unperturbed embryos, despite a lack of individual cell envelopes. These appear to be needed for fine-tuning the chronology of sequential division events, hence allowing all cells to relax to the desired loci in due time.

Going beyond the mere timing, we next asked whether space compartmentalization in the syncytium still follows the cell arrangement observed in unperturbed embryos. Given that cells had been seen to relax into positions of least constraints within the ellipsoidal eggshell [14, 15], this would mean that the relative positioning of syncytial nuclei is similar to those of cells in the unperturbed embryo. For a proper comparison between unperturbed and squeezed embryos, we have labeled nuclei in the syncytium of compressed embryos in accordance with the corresponding cell names in unperturbed embryos. To this end, we first defined the anterior-posterior axis (AP axis). Since the syncytium retained the ellipsoidal shape of unperturbed embryos, we have set the longest axis as the AP axis and defined the anterior pole to be the one in which at least one of the polar bodies remained immobilized (Fig. 5a). This definition is in full analogy to key features of unperturbed embryos [34]. Nuclei of the first generation in the syncytium were then identified with the somatic AB cell (anterior) and the germline precursor P1 (posterior). For the second generation in the syncytium, daughter nuclei of the AB nucleus were labeled as ABa and ABp, depending on whether they were more anterior or posterior (see Fig. 5a). Similarly, the posterior nucleus emerging from division of the P1 nucleus was labeled as P2, the second daughter nucleus was then named EMS. This assignment of labels becomes more complicated from the third generation onwards since the left-right body axis might be needed to distinguish descendants of nuclei. For simplicity, we have therefore restricted our analysis of the mutual distances of nuclei to the first two generations in which a proper labeling was easily possible.

**FIG. 5.**
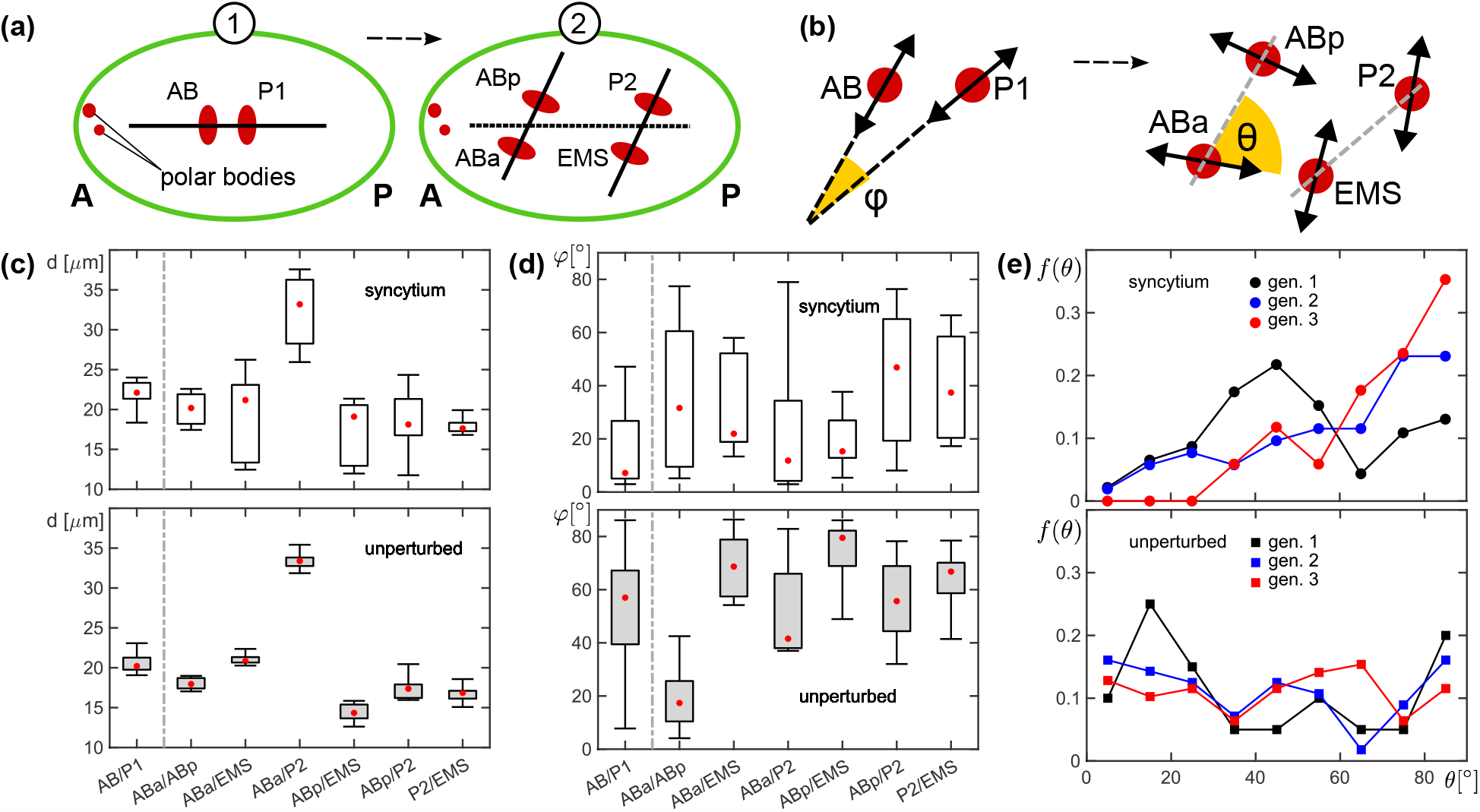
(a) Sketch of how the AP-axis and labels for nuclei were assigned in the syncytium state; see main text for details. (b) Definitions of the angle *φ* between two division axes of nuclei within the same generation and of the angle *θ* between successive division axes (mother/daughter). (c,d) Distances *d* and relative angles of division axes *φ* for the named pairs of nuclei in the first and second generation. While the distances are very similar in unperturbed embryos (bottom) and in the syncytium (top), the orientation of division axes is markedly different. (e) Relative probabilities, *f* (*θ*), for the angle between mother and daughter nuclei. Named generations refer to daughter nuclei, i.e. gen. 1 refers to *θ* values of AB and P1 relative to the AP axis (the division axis of P0). For unperturbed embryos, no clear trend for a preferred angle is visible (except a peak at low angles in the first generation). Successive divisions in the syncytium, however, develop an increasing preference for orthogonal orientations as the generation number increases. See main text for details and discussion.

As a result, we observed that the mutual distances of nuclei in the syncytium follow the cell/nucleus positioning in unperturbed cells (Fig. 5c). In the first generation, the distance between AB and P1 nuclei is basically the same in the syncytium and in unperturbed embryos. In the second generation, stronger fluctuations are observed but the distances are on average still maintained in the syncytium. In fact, the four-cell state of unperturbed embryos, corresponding to the second generation, had been seen to feature a (coplanar) diamond-shaped arrangement of cells/nuclei with the longest distance between ABa and P2 [14] while all other mutual distances were quite similar. This feature is also seen in the syncytium. These data suggest that nuclei in the non-natural syncytium state also follow an optimization criterion for their spatial arrangement, namely to assume positions of least mechanical constraints in an ellipsoid, similar to what has been seen before in unperturbed embryos.

Interestingly, the relative angles of division axes, *ϕ*, within the first and second generation (see Fig. 5b for a defining sketch) deviate considerably between syncytium and native embryos. In unperturbed embryos, the division axes of AB and P1 enclose an angle of about 45° whereas the corresponding nuclei in the syncytium divide considerably more collinear (see Fig. 5d). Similar observations have been made in explants of the syncytial blastoderm of *D. melanogaster* with only two dividing nuclei [31]. Therefore, the diamond-shaped arrangement of second-generation nuclei in the syncytium is not already predetermined by the division axes of first-generation nuclei but rather appears to be driven subsequently by repulsive forces, e.g. mediated by microtubule asters that push nuclei to their final positions [31].

Relative angles *φ* between division axes of syncytial nuclei of the second generation also deviate from the unperturbed case. While in unperturbed embryos a small angle between division axes is only seen for ABa and ABp (*φ* ≈ 20° instead of *φ* ≈ 60° for all other angles), division axes in the syncytium mostly are in the range of small angles (*φ* ≈ 20°), albeit with very strong fluctuations. These observations suggest that a proper orientation of division axes requires functional cells, i.e. a plasma membrane that can host guiding cues like PAR protein gradients or (chiral) actomyosin forces in the cell cortex. In other words, without a cell-defining membrane envelope, the spatial arrangement of nuclei can still resemble the unperturbed case in a robust fashion but the well-defined orientations of division axes of unperturbed embryos are basically lost, resulting in a preference for more collinear divisions. Interestingly, the latter finding deviates from the aformentioned explants of syncytial blastoderms of *D. melanogaster* with more than two nuclei, for which a successive randomization of the angle *φ* has been observed [31]. This difference might arise from the fact that nuclei in the syncytium of *C. elegans* embryos are still confined by a rigid eggshell that can provide mechanical cues and boundary conditions, whereas nuclei in explants of the fly embryo are virtually unconfined.

To inspect the geometry of divisions in some more detail, we also extracted the angle enclosed between successive divisions (see sketch in Fig. 5b). In fact, comparing the angle *θ* between a nucleus division axis in generation *n* and the division axes of its two daughter nuclei in generation *n* + 1 reports on memory effects in successive divisions and whether these necessitate a cell membrane. As a result of our measurements, we observed that nuclei in unperturbed embryos feature a broad range of division angles in each generation with little preference for a particular value of *θ* (Fig. 5e). Only in the first generation, a distinct peak at small angles is seen that captures the division of P1 preferentially along the AP-axis (the division axis of the mother cell P0). From the second generation onwards, there is no clear preference for a distinct angle between successive divisions, indicating that selecting division axes is basically devoid of a memory in unperturbed embryos on the level of the whole cell ensemble (individual cells like P1 or EMS still show a preference for certain angles). In contrast, the syncytium successively develops a clear preference for a 90° angle between successive divisions for increasing generation number on the ensemble level (Fig. 5e). Please note that here no assignment of cell labels is necessary, i.e. data beyond the second generation were analyzable for syncytia in a meaningful way as well.

The emerging preference for successive divisions with an orthogonal orientation in the syncytium may be rationalized as follows: During anaphase of a division event the two centrosomes are located on opposing faces of the newly forming envelopes of the two daughter nuclei. Using the center of mass of the two emerging daughter nuclei as coordinate center, the two centrosome position vectors r_1_ and r_2_ are, by definition, on the division axis, but point into opposite directions. If the two centrosomes do not move wildly during the following interphase, they will duplicate before the next mitosis event almost at their initial positions, yielding two sister centrosomes per nucleus near to positions r_1_ and r_2_. Assuming each pair of sister centrosomes to interact in a repulsive manner, e.g. mediated by microtubules that originate from these two organizing centers, the two centrosomes will push each other to a position of largest distance on the (circular) nuclear envelope. Consequently, each of the two sister centrosomes will migrate on the nuclear envelope by an angle of about 90°, eventually resulting in the next division axis to be almost perpendicular to the previous division axis, in line with our experimental observation. Therefore, keeping cells in *C. elegans* embryos intact can be seen as a means to prevent a falling back to the default memory that pushes for orthogonal orientations of the mitotic axes of mother and daughter nuclei.

Altogether, our data show that determining the orientation of division axes appears to be tightly guided by individual cells in unperturbed embryos, hence safeguarding the early development. Although this tuning of division axes is lost in the syncytium, the spatial arrangement of nuclei up to the four-nuclei state is still surprisingly similar to what is seen in unperturbed embryos. Certainly, increasing fluctuations in division angles, as well as the successive mixing of chromatids due to a lack of cell membranes, eventually will mess up the spatial arrangement in the non-natural syncytium of mechanically perturbed *C. elegans* embryos, leading to a rapid abort of the development.

## IV. CONCLUSION

In summary, we have shown here that a gentle compression preserves the timing and sequence of cell divisions in *C. elegans* embryos, including a lineage-specific anti-correlation of cell lifetimes and volumes according to Eq. (1). Only upon inducing photoxic damage, periods between successive divisions were significantly increased and the lineage-specfific features were lost, whereas the overall anti-correlation with cell volumes were retained. When compressing embryos to about half of their diameter, cytokinesis was frequently seen to fail, resulting in a non-natural syncytium state that still allows for multiple division cycles of nuclei. The lifetime of syncytial nuclei increases with every generation, in accordance with unperturbed embryos, underlining the robustness of the embryonic timer in early developmental stages. Within the first generations the spatial arrangement of syncytial nuclei is also close to the corresponding cell pattern in unperturbed embryos, suggesting that both are mostly determined by a simple optimization process that favors positions of least mechanical constraints within the ellipsoidal eggshell. In contrast to unperturbed embryos, however, nuclei in the syncytium divided synchronously and displayed altered relative orientations of their division axes, both being reminiscent of observations in the syncytial blastoderm of *D. melanogaster*).

Altogether, our findings suggest that few robust physico-chemical cues provide a basic developmental program for timing mitotic events and for guiding the subsequent embryonic compartmentalization, allowing the early development of *C. elegans* to run virtually on autopilot.

## AUTHOR CONTRIBUTIONS

VB performed and analyzed all experiments and added to the writing of the manuscript; MW designed research, complemented the data analysis, and wrote the manuscript.

## ACKNOWLEDGMENTS

Financial support by the DFG (grant WE4335/3-2) and by the VolkswagenStiftung (Az. 92738) are gratefully acknowledged. Some strains were provided by the Caenorhabditis Genetics Center funded by the NIH Office of Research Infrastructure Programs (P40 OD010440).

