## Supplement for "Robust spatiotemporal organization of mitotic events in mechanically perturbed *C. elegans* embryos"

### Figure S1

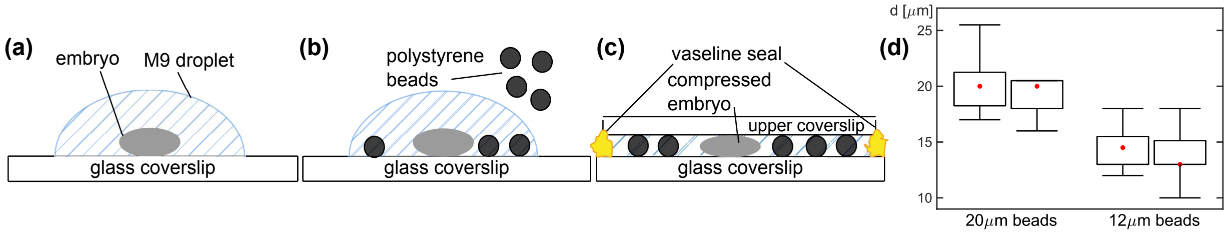


FIG. S1 Illustration of the protocol to compress *C. elegans* embryos. (a) The embryo is placed inside a droplet of M9 buffer on a glass coverslip. (b) Then a solution of M9 buffer, mixed with polystyrene beads of the desired diameter, is added to the droplet. (c) Finally, a second coverslip is placed on top of the droplet, resting on the polystyrene beads, hence compressing the embryo. Sides are sealed by gently applying hot Vaseline to prevent movement and medium evaporation. (d) The optically determined distance $d$ between the two coverslips (when using 20 $\mu$m and 12 $\mu$m spacer beads) provides information on the thickness of the embryo. Values for $d$ did not vary significantly between the beginning and the end of embryonic time series (left vs. right box plots). We attribute the slightly higher values for $d$ when using 12 $\mu$m beads to variations of bead diameters (measured to cover the range $11.5 - 13.5$ $\mu$m) and, to a minor extent, to the elastic response of strongly squeezed embryos.

### Figure S2

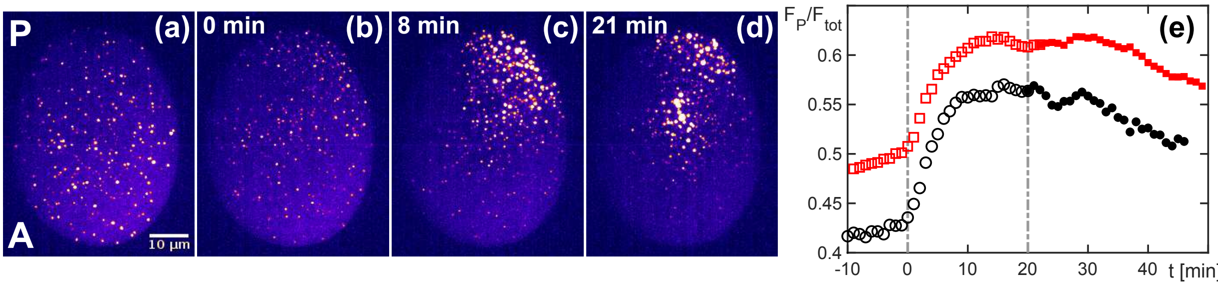


FIG. S2 Despite a compression of the embryo that prevents cytokinesis, p-granules still form and accumulate at the posterior end of the embryo before the first mitosis event. A roughly equi-distributed pattern of p-granules is seen (a) about 10 min prior to and (b) at the time of the first division of nuclei (defined here as $t = 0$ min). (c) Shortly after the first division, a proper spatial gradient has emerged, similar to what is seen in unperturbed embryos before the first cell division. (d) At and beyond the second division of nuclei, the gradient pattern gradually subsides. Instead, p-granules accumulate near to dividing nuclei in the center of the syncytium. (e) The fluorescence fraction in the posterior half of the embryo, $F_P/F_{texttot}$ reflects an enrichment of p-granules at and beyond the time of the first division event. Please note that the fluorescence also includes the cytosolic background fluorescence of unbound gpl-proteins.

*Contact: ``
